# Clearwing butterflies challenge the thermal melanism hypothesis

**DOI:** 10.1101/2023.07.31.550889

**Authors:** Violaine Ossola, Fabien Pottier, Charline Pinna, Katia Bougiouri, Aurélie Tournié, Anne Michelin, Christine Andraud, Doris Gomez, Marianne Elias

## Abstract

In contrast to most butterflies harboring opaque wing colorations, some species display large transparent patches on their wings. Wing transparency, which entails a dramatic reduction of pigmentation, raises the question of potential costs for vital functions, such as thermoregulation, especially along climatic gradients. The thermal melanism hypothesis posits that darker colorations should be favored in colder environments, which enables them to absorb more radiation and maintain a body temperature compatible with activity. This prediction extends to the near infrared (NIR) range, which represents a large proportion of solar radiation. Here we assess the implications of wing transparency for light absorption and thermal properties in 42 butterfly species from the neotropical tribe Ithomiini that range the extent of transparency, from fully opaque to highly transparent, and we test whether those species conform to the prediction of the thermal melanism hypothesis. We find that transparent wings are less efficient than opaque wings to absorb light across UV, Visible and NIR wavelength ranges, and are also less efficient to collect heat. Moreover, dark coloration occupies a lower proportion of wing area as altitude increases, and ithomiine species harbor more transparency at higher altitudes, where climatic conditions are colder, going strongly against the prediction of the thermal melanism hypothesis. We discuss these surprising results in light of recent studies suggesting that factors other than adaptation to cold, such as predation pressure, physiology or behavior, may have driven the evolution of wing patterns in Ithomiini.

**Significance Statement:** The thermal melanism hypothesis predicts that organisms should be darker and absorb solar radiation more efficiently in colder environments. The Neotropical butterflies Ithomiini are unusual in that many species harbor large transparent patches on their wings, raising questions related to their efficacy of solar radiation absorption and heating capacities. We investigate optical and thermal properties of several ithomiine species along a climatic gradient. We find that transparent wings are less efficient at absorbing radiation and collecting heat. Unexpectedly, the proportion of transparent species increases with altitude, challenging the thermal melanism hypothesis and suggesting that factors other than adaptation to cold, such as predation pressure, may have driven the evolution of wing patterns in Ithomiini.

## Introduction

Diurnal butterflies are known for the great variety of colors they display. The scales covering their wings produce colorations through pigments (1) or structures (2, 3). Coloration is involved in multiple functions, including intraspecific signaling (4); antipredator defenses such as crypsis and camouflage (5), masquerade (6), aposematism and mimicry (7); as well as thermal adaptation (8). The balance between these selective pressures drives the diversity of color patterns in butterfly wings (9).

In ectotherms, coloration has been shown to play an important role in determining body temperature and activity, since the degree of darkness is positively correlated with internal body temperature (10). Darker butterfly wings absorb more light, and are associated with longer flight activity and larger flight distances in cold conditions than lighter wings (11, 12).

The thermal melanism hypothesis (TMH) posits that darker colorations provide thermal advantages in cold environments (13), especially for ectotherms, as dark surfaces are more efficient to absorb heat from the environment than light surfaces. Darker species or individuals are expected to live at high altitudes or latitudes, whereas lighter ones are more likely to be present at lower altitudes and latitudes, which offer warmer conditions. The existence of such melanism gradients has been shown in many case studies, across geographical regions and wide ranges of taxa, from reptiles to insects (14–16). However, visible colors represent only a small part of the incident solar radiation reaching the earth’s surface. Over 50% of the solar energy lies in the near-infrared wavelengths (NIR, 700 to 2500 nm). NIR should then be of particular importance in ectotherm thermal capacities, more than ultraviolet to red (i. e. 300 to 700 nm, hereafter UV-Visible range) (17), since they are not involved in other aspects of their ecology because they cannot be seen by most organisms. Recent studies on communities of European (18) and Australian (19) butterflies confirmed that reflectance in the NIR range is strongly correlated with climatic factors, especially temperature, and more strongly compared to UV and Visible wavelengths. In colder environments, butterflies tend to reflect less and therefore absorb more in the NIR. These findings highlight that optical properties can be disconnected to a certain extent between NIR and UV-Visible ranges. By contrast, no clear trends exist regarding humidity or precipitation and wing darkness. Gloger’s rule, which predicts that darker individual should be found in more humid climates, has been evidenced mainly for endotherms with little clear support in ectotherms (18, 20).

In Lepidoptera, few species exhibit transparency on their wings, a feature that has evolved multiple times independently (21). Transparency is achieved by a large diversity of wing microstructures, including a combination of scale shape or insertion modification, reduced scale size or density, as well as a reduction of scale pigments (21). In many cases, transparency is enhanced by the presence of nanostructures on the wing membrane, which decreases reflection (22–24). Transparency has attracted recent research effort regarding its now-established role in crypsis in various species (25–27), and its involvement in predator-prey signaling in some clades of mimetic Lepidoptera (22). Yet, transparency may come at the cost of other functions involving wing coloration, notably thermoregulation. While thermoregulatory processes are well documented in opaque species, they are largely unknown in clearwing butterflies. A study focusing on wing transparency across the entire Lepidoptera order found that clearwing species tend to transmit less light, and thus absorb more, in high latitudes than in tropical climates, suggesting a thermal cost for butterflies (21). Yet, the actual impact of transparent wing patches on wing thermal properties needs to be investigated.

Among Lepidoptera, Ithomiini is a remarkable neotropical tribe of ca. 400 species (28, 29), 80% of which exhibit transparency on their wings to some extent. Species are highly segregated along the elevational gradient, from warm lowlands up to colder highlands, as high as 3,000 meters (29, 30). Because wing transparency in Ithomiini is achieved by a reduction in membrane coverage by scales (21, 22), and is therefore associated with a lower quantity of pigments, one might expect those areas to poorly absorb solar light. Previous studies of geographical distribution of Ithomiini wing patterns revealed that the proportion of clearwing species increases with altitude, to the extent that all species above 2000 m have transparent wings (29, 30), contrary to the predictions of the TMH. Yet, this observation is based on the UV-Visible range (22, 29, 30), and optical transparency shown in that range may be dissociated from the absorption properties in the NIR.

Here, we investigate for the first time wing traits relevant for thermal adaptation in 42 transparent and opaque Ithomiini species that represent the diversity of wing colorations, by using Infrared imaging and spectrophotometry. We assess the distribution of these wing traits in relation to the climatic conditions the species are exposed to in a phylogenetic framework. Specifically, we characterize and quantify the optical and thermal properties of transparent and opaque wing patches to investigate two mutually exclusive scenarios:

i. *Absorption compensation* :in this scenario, the higher transmission and lower absorption of wavelengths in the UV-Visible range of transparent wing areas are compensated by a high absorption in the NIR, and the thermal capacities of transparent wings may be similar to or even better than those of opaque wings. Under this hypothesis, adaptation to climatic conditions may drive the altitudinal distribution of wing patterns in Ithomiini.
ii. *Thermal cost* :in this scenario, the disconnection between wavelength ranges is not significant enough, with transparent wings transmitting more and absorbing less in the NIR range than opaque wings, therefore, reflecting similar absorption patterns at shorter wavelengths. This results in transparency being costly in terms of intrinsic heating capacities. Under the thermal cost scenario, thermal properties may not be the main target of selection with regard to the altitudinal distribution of wing patterns in Ithomiini.

In addition, assuming that opaque patches - even if not covering a high proportion of the wing surface in transparent species – may follow the thermal melanism hypothesis, we tested whether they represent a higher proportion of wing surface in species found in colder climates, a mechanism that could also compensate for poorer thermal capacities of transparency under the *thermal cost* scenario. The absence of evidence of compensation mechanisms (disconnection NIR from UV-Visible range, or increased proportion of opaque patches with altitude) would imply that factors other than wing color patterns, such as behavior or metabolism, explain adaptation to colder highland habitats in Ithomiini.

## Results

To assess the thermal adaptation of Ithomiini butterflies, we examined the thermal and optical properties of the wings of 42 species in relation to climatic conditions. Unless specified otherwise, all statistical tests implemented Bayesian mixed models with Markov Chain Monte Carlo correcting for phylogenetic relatedness, with an associated Bayesian posterior P-value set to 0.05.

### Wing optical and thermal properties

We first tested whether optical properties in the NIR wavelengths were correlated with those properties in the UV-Visible range, and whether those properties differed between transparent and opaque species, as defined hereafter. We measured transmission and reflection spectra on a 5mm-diameter spot on one hindwing and one forewing for each specimen, spanning the UV-Visible (300-700 nm) and NIR (700-2500 nm). We used the physical principle stating that Transmission (λ) + Reflection (λ) + Absorption (λ) = 1 to compute the total absorption spectra over 300-2500 nm (Fig. 4C). The transmission of the patch in the UV-Visible range was set as a measure for the degree of optical transparency. We split the species in two categories: transparent, for which wing transmission was higher than 35%, and opaque for the others (Table S1, Fig.S1).

Overall, UV-Visible transmission (300-700 nm), i.e., transparency as perceivable by butterflies and predators, was a good predictor of both overall absorption (300-2500 nm) and NIR absorption (700-2500 nm, Table S2, Fig. 1). NIR and UV-Visible absorption were highly correlated (Table S2, Fig. 1), be the species transparent or opaque. More transparent patches absorbed less light than opaque patches whatever the wavelength considered, supporting the *thermal cost* scenario for transparency (Fig. S2).

**Figure 1.**
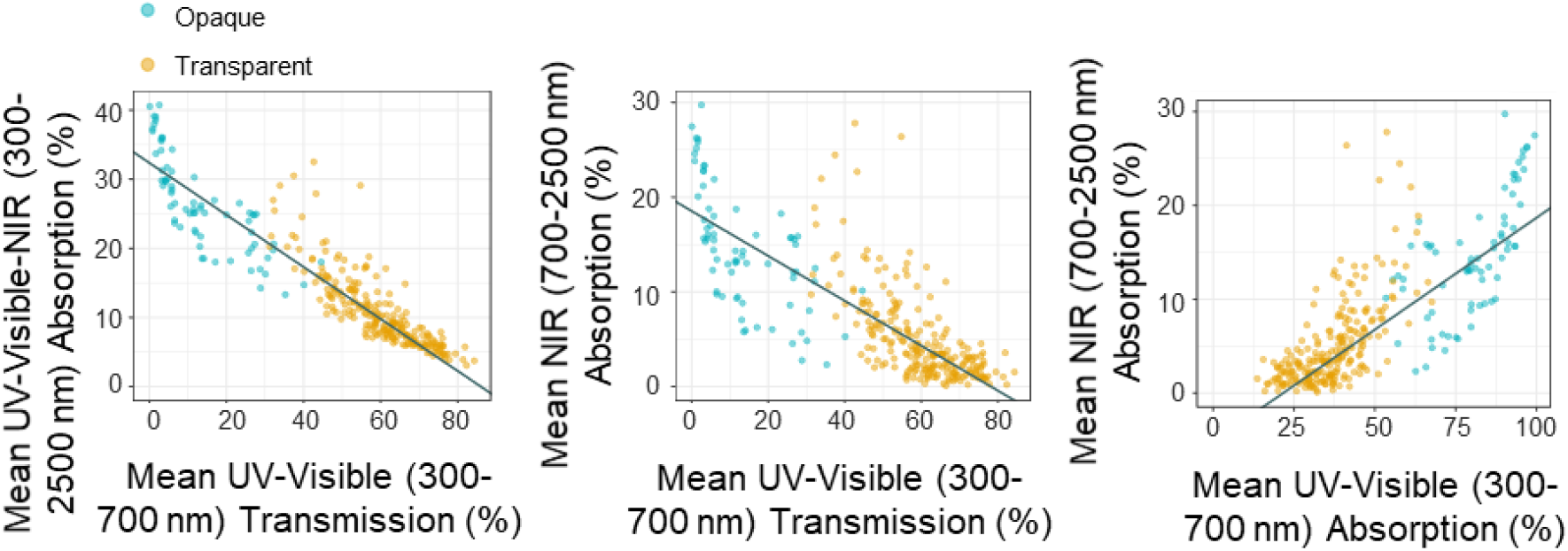
Relationship between absorption and transmission spectra at different ranges: overall UV-Visible-NIR (300-2500 nm) absorption against UV-Visible (300-700 nm) transmission, NIR (700-2500 nm) absorption against UV-Visible transmission, NIR (700-2500 nm) absorption against UV-Visible (300-700 nm) absorption. All relationships are significant and the solid line is drawn from the prediction of the Bayesian phylogenetic mixed model (see Table S2) (N opaque = 72, N transparent = 235).

To investigate the thermal capacities of transparent wings and test whether they differ from those of opaque wings, we heated up wings with a flash illumination, and imaged them with an infrared camera to record temperature across the wing and its variation through time (Fig. 4A-B). Image analysis showed that the average heating capacity of the wing spot, represented by excess temperature ΔT (maximum temperature reached after illumination - initial temperature), was closely related to its transparency, as estimated by the transmission in the UV-Visible range (Table S3, Fig. 2). Hence, the more transparent the spot, the less it heated up, in agreement with the *thermal cost* scenario. ΔT of wing spots was also positively predicted by the mean total absorption of the spot and by its NIR absorption (Table S3, Fig. 2). Therefore, wings that absorbed more in the UV-Visible-NIR range or in the NIR range heated up more. In an in-depth exploration of the intra-wing variation in heating capacity, we extracted ΔT on various points on different parts of the wings (veins and cells, i.e., the parts between veins), either in darkly colored patches DP (black to brown colors), lightly colored patches LP (for opaque species only, yellow to orange colors), or transparent patches TP, and on light LV or dark veins DV. A close look at the different areas and structures of the wings revealed that dark patches heated up significantly more than any other coloration on the wings, transparent or light (DP>(LP,TP) in Table S4, Fig. S3). Moreover, light patches heated up significantly more than transparent patches (LP>TP in Table S5, Fig. S3). Dark patches heated up more than dark veins (DP>DV in Table S6, Fig. S3), but on the other hand, light veins absorbed more heat than light patches (LV>LP in Table S7, Fig. S3). In all cases, the position of the patch with respect to the thorax, proximal or distal, played a role in predicting ΔT, but this varied according to the color of the patch considered, showing no clear-cut trend (Table S4-7).

**Figure 2.**
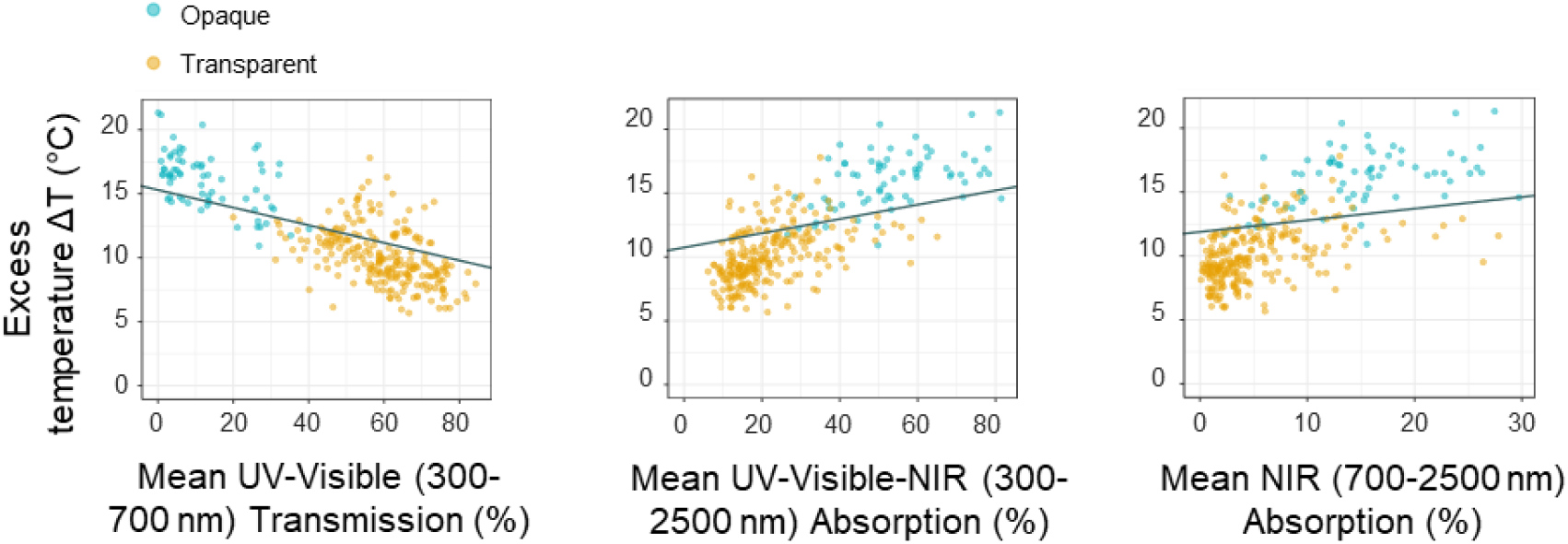
The thermal properties of wing spots represented by the Excess Temperature ΔT (T°final – T°initial) under a flash illumination, extracted from the thermal imaging, against their optical properties at different ranges: UV-Visible (300-700 nm) transmission, UV-Visible-NIR (300-2500 nm) absorption, and NIR (700-2500 nm) absorption. All relationships are significant and the solid line is drawn from the prediction of the Bayesian phylogenetic mixed model (see Table S3) (N opaque = 72, N transparent = 235).

### Wing trait distribution along climatic gradients

We tested whether climatic variables predicted the proportion of dark patches on the wings of all the species, as posited by the TMH. We extracted 19 published average climatic variables for each species (29) that included temperature and precipitation descriptors and performed a PCA. PC1 and PC2 explained respectively 49 % and 27.1 % of the total variation. Higher PC1 values corresponded to warmer climates, while higher PC2 values corresponded mainly to wetter and more time-stable climates (tables S8, S9, Fig. S4). Pearson’s product-moment correlation tests showed that PC1 was strongly negatively correlated with altitude (R=-0.95, N=42, p <.0001), meaning that species inhabiting higher altitude habitats had a lower PC1 value. PC2 did not correlate with altitude (R=0.24, N=42, p=0.12). To test if opaque and transparent species had similar climatic ranges we used the Levene’s heteroscedasticity test on climatic variables. We found that opaque species were restricted to warmer environments, while transparent species were ubiquitous and not associated to particular altitudes (F =110.71, p <.005) (transparent species: mean=-0.27, var=10.56, opaque species: mean=1.08, var=1.09), in contradiction with the TMH. Opaque and transparent species did not significantly differ in their variance for PC2 (F=1.76,p=0.19) (transparent species: mean=0.29, var= 3.88, opaque species: mean=-1.89, var=3.45), showing that they did not segregate according to humidity or environmental stability gradients. Overall, climate did not predict wing optical properties. The degree of transmission of the wing spots over the range UV-Visible were not predicted by PC1, with a negligible positive effect for PC2 which goes against Gloger’s rule (Table S10, Fig. S5). There was no significant association of spot absorption in the UV-Visible-NIR (table S11) or NIR ranges (table S12) with PC1 or PC2. By contrast, the average ΔT of the whole wing surfaces was positively correlated to PC1 (Table S13, Fig. 3). In other words, species living in warm environments heated up more than butterflies living in colder environments.

**Figure 3.**
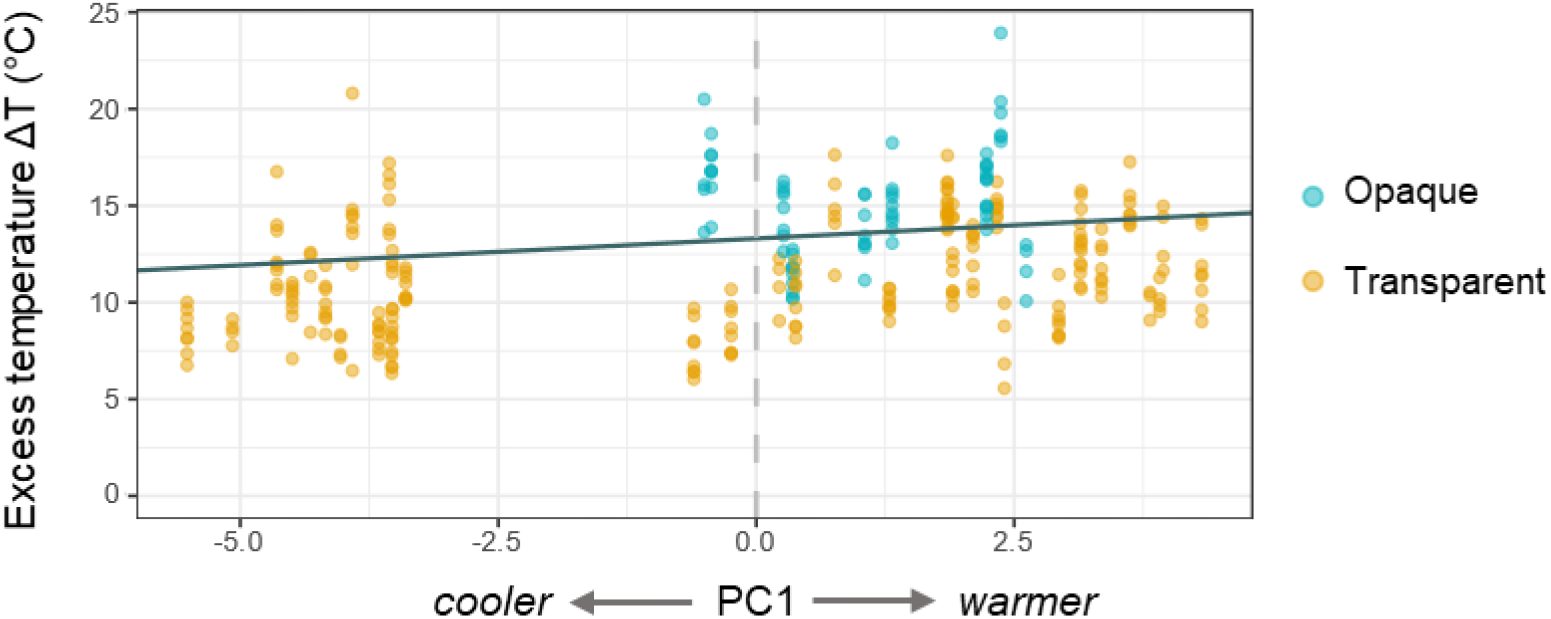
Wings mean Excess Temperature ΔT(T°final – T°initial) under a flash illumination, extracted from the thermal imaging, as a function of climatic principal component PC1. The relationship is significant and the solid line is drawn from the prediction of the Bayesian phylogenetic mixed model (see Table S13) (N opaque = 72, N transparent = 235).

Across all species, the proportion of dark surface on the wings increased with PC1 and was therefore higher in warmer environments (Table S14, Fig. S6). We found a similar trend when restricting the analysis to only transparent species (Table S15, Fig. S7). We examined whether this proportion varied with climate within the proximal zones alone, because wing conduction could be limited, and a patch was expected to have a greater thermal effect when closer to the body. Again, the proportion of dark surface in the proximal zones increased with PC1 (Table S16, Fig. S8), and was therefore higher in warmer environments, in contradiction with the TMH. The proportion of dark surface did not correlate with PC2, except when considering only proximal zones, indicating that proximally darker wings were associated with more humid habitats. Finally, we found no effect of climatic variables on total wing area, a proxy for butterfly size (Table S17).

## Discussion

### Thermal properties correlate with optical properties in clearwing butterflies

The thermal imaging of Ithomiini wings reveals that opaque wings are on average more efficient to heat up than transparent wings. Hence, transparency entails thermal costs, a conclusion supported by the close-up study of the different wing patches, in agreement with the *thermal cost* scenario. The darker the color, the more the patch heats up. Therefore, transparent patches contribute little to wing heat absorption, in line with the long studied link between melanization and heating capacities (31). Consistent with this finding, the optical examination of the transparent wing spots reveals that the low absorption in the UV-Visible wavelength range is not compensated by a higher absorption in the invisible NIR range, and therefore wing transparency comes at the cost of light absorption. This result departs from findings in other organisms where visible reflectance tends to be a poor predictor of NIR reflectance (32). Moreover, transparent spots always absorb less light than opaque ones, regardless of the wavelength considered. This is in agreement with the correlation of the UV–Visible and NIR spectra that has been previously described for transmission(21) or reflection (18, 19) in interspecific studies of butterflies taxa. Yet, veins in transparent Ithomiini species are always covered with melanized scales. Those veins heat up significantly more than transparent and even lightly colored patches, and likely represent a major part of the overall wing thermal capacities by providing heat to the dynamic hemolymph flow system to the thorax (33).

In Ithomiini, absorption decreases in long wavelengths, and is quite low in the NIR for all the studied species. An interspecific comparison of the absorption of longer wavelengths (∼8-12 μm), especially infrared radiated by the environment, remains to be carried out as this wavelength range is supposed to have greater thermal impact. However, the thermal imaging results clearly show that transparent patches are poorer at accumulating heat than opaque patches, suggesting an absence of compensation in the far infrared range.

### The altitudinal distribution of transparent species contradicts the thermal melanism hypothesis

This study confirms the counterintuitive altitudinal distribution of Ithomiini wing patterns (30). While the opaque species are only found in warm climates, clearwing species are ubiquitous and widely distributed, from hot lowlands to the cold Andean highlands. Moreover, among clearwing species, the proportion of opaque patches on the wings decreases in colder environments, i.e. at higher altitudes, challenging the long history of studies supporting the TMH in butterflies and insects e.g.(15, 16, 34, 35), but adding to the observation that TMH does not apply to all taxa or communities. For instance a recent study found support for TMH in geometrids but not in noctuids found in the same study sites (36). Yet, these studies often focus on temperate climates. Neotropical case studies are scarce and show mixed support to the TMH. Two closely related neotropical butterfly genera, Catastica and Leptophobia have a divergent coloration distribution along altitudinal gradients (37), possibly explained by different behaviors and elevation ranges. While body and wing colorations become darker with increasing altitude in Catastica, Leptophobia shows a reversed pattern, similar to that observed in Ithomiini. In the case of Ithomiini, not only the non-conformity to TMH is outstanding, but there is no high NIR absorbance to compensate the poor UV-Visible absorption that would make highland butterflies more adapted to cooler conditions.

The NIR range is expected to be only subjected to thermoregulatory pressures, unlike UV-Visible wavelengths involved in vision. Indeed, NIR reflection spectra of Australian and European butterflies communities are well predicted by climatic conditions (18, 19). In contrast to their conclusions, our study of the absorption properties of wing spots showed that in general, climatic conditions do not predict wing absorption in any radiation range. Moreover, the exploration of wing heating capacities shows that Ithomiini species living in colder environments are less efficient to heat up than species living in warmer climates, adding even more to the thermal cost of transparency. Finally, there is hardly any link between thermal or optical properties and humidity, and as such this study does not support Gloger’s rule in Ithomiini.

Adaptation to cooler conditions is not the only thermal constraint faced by organisms, as high temperature may also be challenging. Here we found that opaque species, which absorb more radiation and have higher thermal capacities, live in environments where they are most exposed to heat. For ectotherms, avoiding overheating (i.e. increasing body temperature above a critical temperature threshold) can be challenging. Butterflies may have physiological adaptations (38) or behavioral adjustments, such as changing the position of their wings according to environmental radiations (33). Information on the metabolism and behavior of Ithomiini *in natura* would shed light on their altitudinal adaptation, but are currently lacking. In addition, microhabitat variability can result in individuals sharing the same overall environment being exposed to contrasting light and temperature conditions. Microhabitat preferences thus likely play a role in adjusting body temperature (38, 39). However, in lowland ithomiine communities, butterflies with transparent wing patterns mostly fly in the understory, close to the vegetation and out of direct sunlight, while the proportion of species with opaque wings increases with height in the canopy (40). The latter are therefore more exposed to solar radiation, another counterintuitive distribution in relationship to thermoregulation.

### Transparency under multiple selection pressures

Colorations in butterflies, and therefore wing optical properties, are under multiple selection pressures as they are involved in anti-predator defenses, signaling to conspecifics and thermal adaptation, with potential trade-offs (9). Consequently, selection for crypsis and aposematism could be detrimental for thermal capacities (41). Wood tiger moths are a good example of the phenomenon: larvae with large black patches are at a thermal advantage compared to larvae with equally black and orange patches, but the latter display a more efficient aposematic signal (42). Imagoes display latitudinal variation in wing pattern, consistent with the TMH, which incurs cost in the efficiency of warning signals as melanic individuals experience more predation (34). In Ithomiini, the existence and pervasiveness of transparency is surprising given that those butterflies are toxic and harbor aposematic signals. Previous work has shown that transparency in Ithomiini reduces detection and attacks by naïve predators (27). Even though transparent patches are likely part of the aposematic signal (22), transparency may reduce predator learning speed and memorization of prey, as indirectly suggested by the observation that unpalatability tends to be stronger in clearwing than in opaque species (43). Thus, transparency may incur a cost on aposematism, with a trade-off between crypsis and aposematism. A possible explanation for the unexpected altitudinal distribution of transparency in Ithomiini species may be different predation pressures at different altitudes in tropical mountains (44), which may favor crypsis over aposematism at higher altitudes, as observed in Andean arctiine moths or which conspicuousness decreases with height (45). Therefore, the altitudinal distribution of color patterns in ithomiine may be primarily driven by predation rather than climatic conditions.

## Conclusion

In Ithomiini, the altitudinal distribution of wing patterns is unlikely driven by thermal constraints, and goes against the predictions of the thermal melanism hypothesis. Transparent wings have a low absorption of wavelengths in the UV-Visible-NIR range and this comes at a cost regarding heating capacities. Conversely, transparency in these butterflies has probably evolved in response to selection incurred by predation. The results question the critical significance of wing color pattern for thermal adaptation in tropical ectotherms and lead the way to further investigations about metabolic and behavioral thermoregulation, and their relative contributions to climatic adaptation in these species.

## Materials and Methods

### Specimen sampling, climatic data and phylogeny

We studied wings of dried Ithomiini butterfly specimens, preserved in glassine envelops, collected between 2005 and 2018 in south America. We focused on 42 species represented by two or more females to reduce the potential variance due to sexual differences (Table S1). Measurements were taken on one hindwing and one forewing of each specimen. The selected species spanned a large range of wing coloration, from fully opaque to highly transparent, and are evenly distributed along the altitudinal gradient and across the phylogeny. GPS data on species distributions were obtained from a database that contains more than 3500 occurrences of specimens of the species studied here (29). We extracted the corresponding 19 climate variables (Table S8-9., Fig. S4) from WorldClim2 historical climatic data (46) with a resolution of 2.5 arc-minutes. Variables were averaged for each species. For the phylogenetic comparative analyses, we used the most comprehensive phylogeny of the Ithomiini to date (28), which comprises 340 out of 393 extant taxa.

### Spectrophotometry measurements

To quantify the extent to which wings transmit and absorb light, we performed spectrophotometry measurements on a spot of 5 mm of diameter on the hindwing and forewing of each specimen. For transparent species, the spot was in the proximal transparent patch. For opaque species, the spot was located on the center of the proximal zone of the wing. Since a wing can reflect and transmit a large proportion of the incident light, we measured both total (specular + diffuse) reflection and total (specular + diffuse) transmission spectra of each patch, with an Agilent CARY 5000 UV-Visible-NIR spectrophotometer, equipped with an integrating sphere, over the range 300-2500 nm, which includes ultraviolet (UV), Visible, and near-infrared (NIR) wavelengths. Wings were maintained in a black paper mask with an opening of 5 mm of diameter. Spectra were measured relative to a dark (paper mask without the wings, light off) and to a white reference (Spectralon white diffuse reflectance standard), with the paper mask for reflection, nothing for transmission, light on. The resulting total absorption spectra were calculated for each wavelength λ as Absorption (λ) = 1 - (Transmission (λ) + Reflection (λ)) according to the law of conservation of energy (Fig. 4). The negative values due to the noise of the signal were fixed to zero with the function fixneg of the package Pavo in R (47), and each spectrum was smoothed with the loess function in R. We quantified mean absorption at different wavelength ranges, in the UV-Visible between 300-700 nm, in the NIR between 700-2500 nm, and the total 300-2500 nm. Mean transmission in the UV-Visible range 300-700 nm was measured as a proxy for animal-visible transparency. We divided the species into two categories, transparent and opaque, based on a threshold of mean transmission at 35% in the UV-Visible, matching butterflies and predators’ visual perception (Table S16).

**Figure 4.**
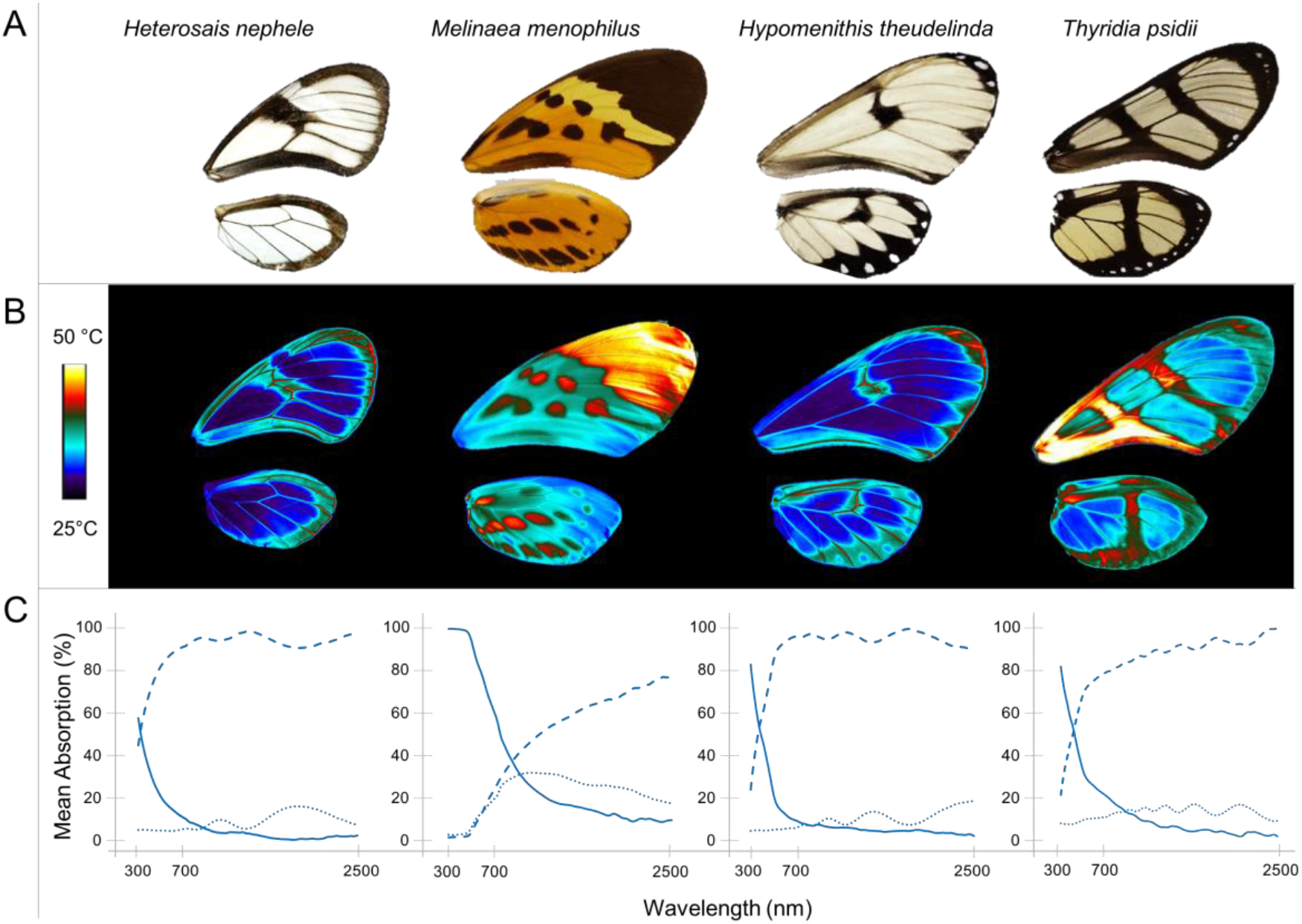
Examples of optical and thermal imaging on the wings of four species of different opacity. A: Photos under standard illumination of the hindwings and forewings of four of the species studied, *Heterosais nephele, Melinaea menophilus, Hypomenithis theudelinda, Thyridia psidii*. B: Visualization of the thermal imaging in false colors of the same wings under a flash light. C: Corresponding measured transmission (dashed lines) and reflection (dotted lines) spectra in percentage, for a spot of 5 mm of diameter on the proximal forewing of each butterfly, from 300 to 2500 nm, and the resulting computed absorption spectra (solid lines) for each wavelength λ as Absorption(λ)+ Transmission(λ) + Reflection(λ) = 1.

### Thermal measurements

To characterize and compare the wing thermal characteristics, we measured wing temperature kinetics following instant exposure to a flash light (Fig. 4B). We conducted stimulated heating measurements on one hindwing and one forewing for each specimen, with a Neewer professional photo system flash lamp N-300W set at maximum power, which emits over the UV-Visible-IR range. Wings were recorded by an infrared camera T650sc 2019 FLIR Systems equipped with a FLIR lens T197-915. Emissivity was set at 0.95. Wings were kept at a constant ambient temperature of 23°C prior to the experiment. We placed each single wing horizontally on a custom grid made of 3 human hair in a frame to minimize contact with any surface, 6 cm below the camera and the flash and 30 cm above a table covered with white paper, to avoid background interference with the measure. The flash lasted for 1/800 seconds. We started the recording a few seconds before activating the flash, and let the record run for 10 seconds. Each movie was recorded at 30 images.second^-1^. From the movies, we extracted the thermal kinetics - temperature as a function of time - of different regions of interest on the wings with FLIR ResearchIR Max software, and more specifically the excess temperature ΔT as the difference between the maximal temperature attained by the region of interest after the flash hits minus the temperature prior to the flash. We extracted mean ΔT of the overall wing surface, mean ΔT of the spot measured in spectrophotometry and finally ΔT of precise points of interest on each wings as follows: 6 points in the dark colored patches DP (black to brown colors), 6 points in light colored patches LP (yellow to orange colors and for opaque species only), 6 points in the transparent patches TP (for transparent species only), and 8 points on veins, assigned to a category depending on their color: dark veins DV (black to brown) and light veins LV (yellow to orange).

### Image analysis

We used digital photography to quantify the proportion of dark pigmented surface on the full wings of all the specimens. Melanization in proximal zones of the wing can be of primarily importance in thermoregulation as the zone is closer to the thoracic muscles (8). We also quantified the color proportion separately on the proximal and distal zone of each wing. We set the limit between the zones as the perpendicular line to the middle of the transect line from the wing insertion to its apex. We cropped the photos for each zone considered and the background was filled with green, a color ignored when calculating the color proportions. In R, we used the getKMeanColors function from colordistance package (48) to sort the pixels in clusters according to their RGB values. We set the number of clusters to two, corresponding to mainly dark and brown, versus transparent, or any other light colors in opaque wings. With ImageJ 1.52 (49) we computed the total surface of each wing in mm^2^.

### Statistical analyses

We explored various scenarios linking absorption, transmission, thermal capacities and climatic variables across the 42 species of the dataset (see supplementary materials for model details and results). Because specimens are not independent and the species are related through the phylogeny, we implemented for each test Bayesian mixed models with Markov Chain Monte Carlo, correcting for phylogenetic relatedness, using the R package MCMCglmm (50) and the phylogeny from (28). We controlled for chain convergence visually and with the Heidelberg stationarity test diagnostic function of convergence heidel.diag from the R package coda (51). To get accurate estimations of posterior distributions and avoid chain autocorrelation, we adjusted the iterations, burn-in and thinning for each model to ensure an effective sample size across all parameters of at least 1000. Models were run with a weakly informative prior for random effect and residual variances (V = 1, nu = 0.002) not to constrain the exploration of parameter values. We selected the best model with a backward selection of fixed parameters based on Bayesian P-value of 0.05, within the 95% confidence interval excluding zero. In all the models, species identity was implemented as a random factor. We included all individuals in the models unless specified otherwise.

## Acknowledgments

This work was funded by Clearwing ANR project (ANR-16-CE02-0012), HFSP project on transparency (RGP0014/2016). VO is recipient of a doctoral fellowship from the doctoral school FIRE, Learning Planet Institute. We are strongly grateful to authorities in Peru and Ecuador for providing collecting permits (Ecuador: 005-IC-FAU-DNBAPVS/MA, 019-IC-FAU-DNBAPVS/MA021; Peru: 236-2012-AG-DGFFS-DGEFFS, 201-2013-MINAGRI-DGFFS/DGEFFS, 002-2015-SERFOR-DGGSPFFS, 373-2017-SERFOR-DGGSPFFS; Gobierno Regional San Martín PEHCBM: 124-2016-GRSM/PEHCBM-DMA/EII-ANP/JARR). We warmly thank Raúl Aldaz, Paula Santacruz and Alexandre Toporov for providing help in the field, and Santiago Villamarin, the Museo de Historia Natural and Professor Gerardo Lamas for their support. We also thank Jean-Luc Bodnar and Kamel Mouhoubi from Reims University for technical advice with IR imaging, Pierre de Villemereuil for his help with Bayesian analysis, Maël Doré for bioclimatic and distribution data sharing, Willy Daney de Marcillac for his advice with optical measurements and Serge Berthier for inspiring discussion on clearwing butterflies.

