## Supplementary material for "Clearwing butterflies challenge the thermal melanism hypothesis": SI_Figures_S1-S8_Tables_S1-S17

**This PDF file includes:**

Figures S1 to S8
Tables S1 to S17

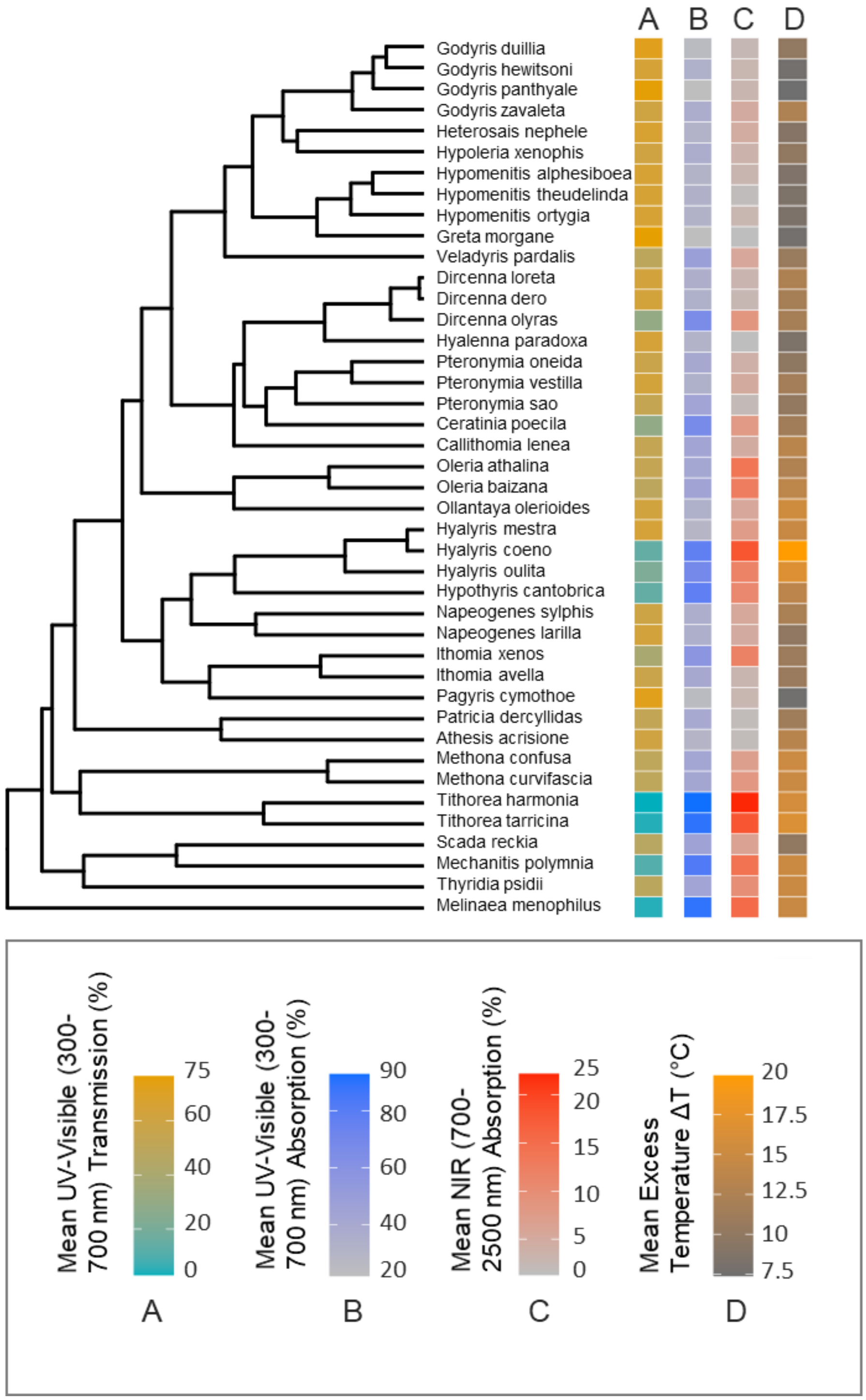


**Figure S1.** Distribution of optical and thermal traits along the phylogeny of the 42 ithomiine species studied, averaged for both wings and all the specimens of each species. A: Mean UV-Visible (300 to 700nm) transmission for a spot of 5 mm of diameter on wing proximal zone. This value is used as a proxy for wing transparency. B: Mean UV-Visible (300 to 700 nm) absorption for a spot of 5 mm of diameter on wing proximal zone. C: Mean NIR (700 to 2500 nm) absorption for a spot of 5 mm of diameter on wing proximal zone. D: Mean Excess Temperature ΔT (T°final – T°initial) in °C of the wings under a flash illumination.

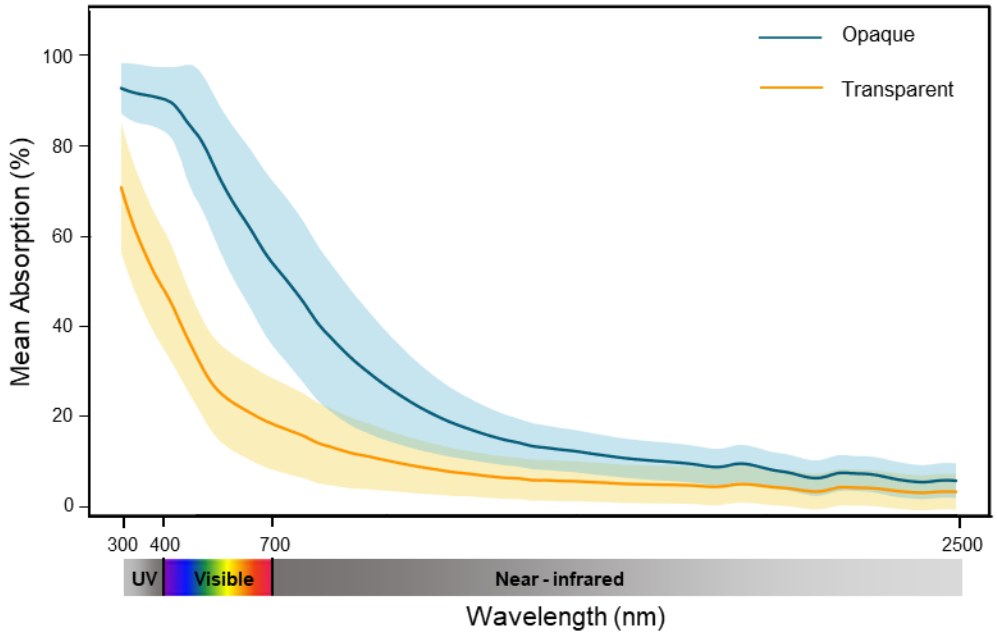


**Figure S2**. Mean absorption spectra and 95% standard deviation of opaque and transparent wing spots from 300 to 2500 nm. Absorption over the whole spectrum is significantly correlated with transmission in the UV-Visible range, as tested by the Bayesian phylogenetic mixed model (see Table S2) (N opaque species = 9, N transparent species = 33).

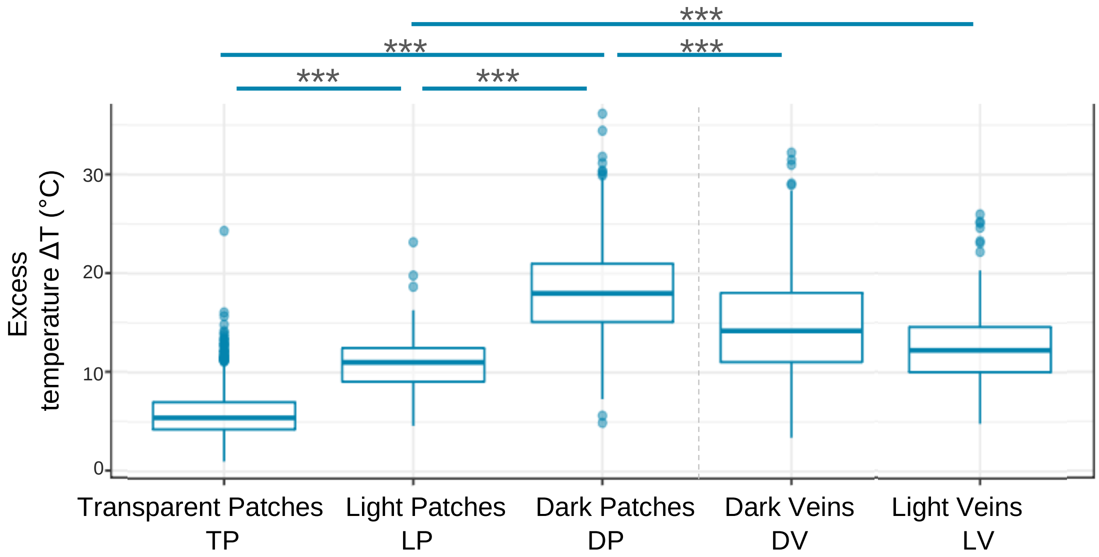


**Figure S3.** Excess Temperature ΔT (T°final – T°initial) of different color patches and veins on the wings, under a flash illumination, extracted from the thermal imaging. The tested pairs shown here correspond to the Bayesian phylogenetic mixed model in the tables S4 to S7. The significance of the effects correspond to an estimated Bayesian P-value (*P < 0.05; **P < 0.01; ***P < 0.001) (N TP= 1407, N LP= 408, N DP=1829, N DV=2117, N LV= 325).

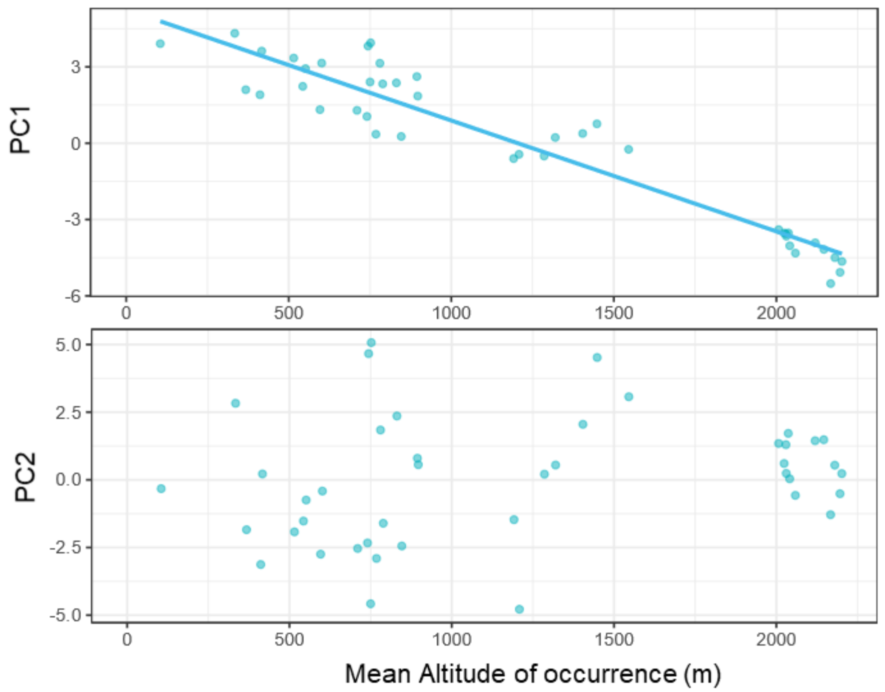


**Figure S4**. Predicted coordinates of the studied species on the climatic principal components as a function of their mean altitude of occurrence. The relationship is significant for PC1 (Pearson's product-moment correlation test: R= -0.95, n=42, p < .0001) but not for PC2 (Pearson's product-moment correlation test: R=0.24, n=42, p=0.12) (N = 42).

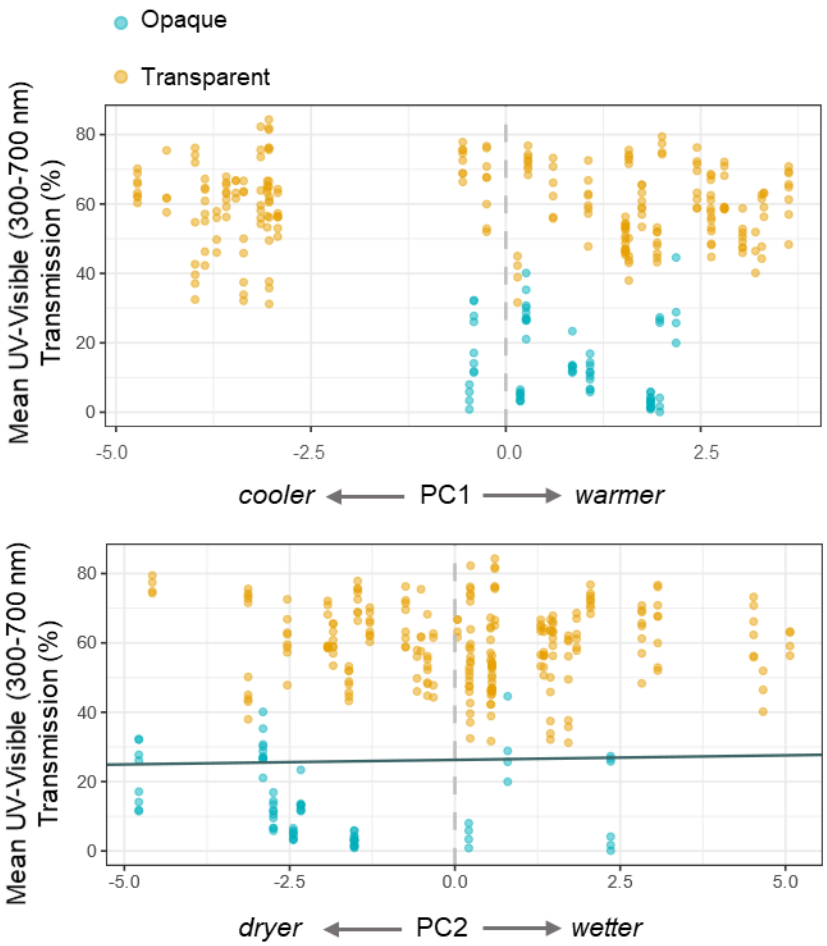


**Figure S5.** Wing mean transmission (%) in the UV-Visible range (300-700 nm) as a function of climatic principal components PC1 and PC2. The relationship is not significant for PC1, and is significant for PC2 (see Table S10), and the solid line is drawn from the prediction of the Bayesian phylogenetic mixed model (N opaque = 72, N transparent = 235).

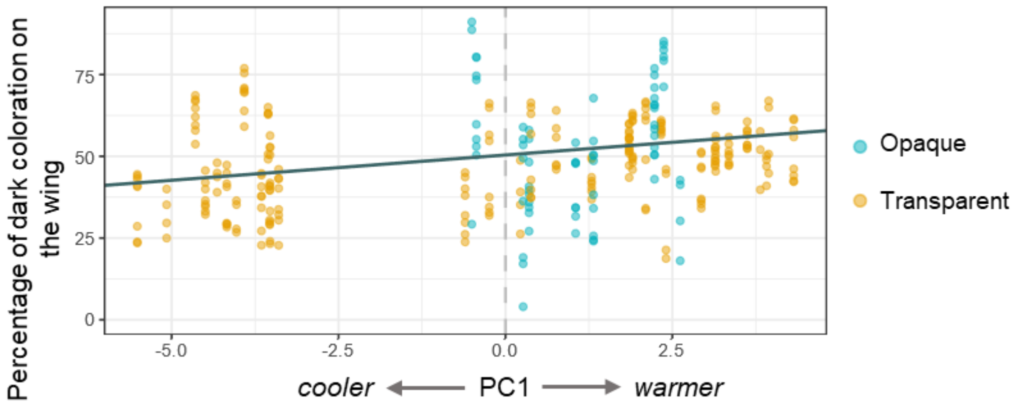


**Figure S6.** Proportion of dark coloration on the surface of butterfly wings as a function of climatic principal component PC1. The relationship is significant and the solid line is drawn from the prediction of the Bayesian phylogenetic mixed model (see Table S14) (N opaque = 72, N transparent = 235).

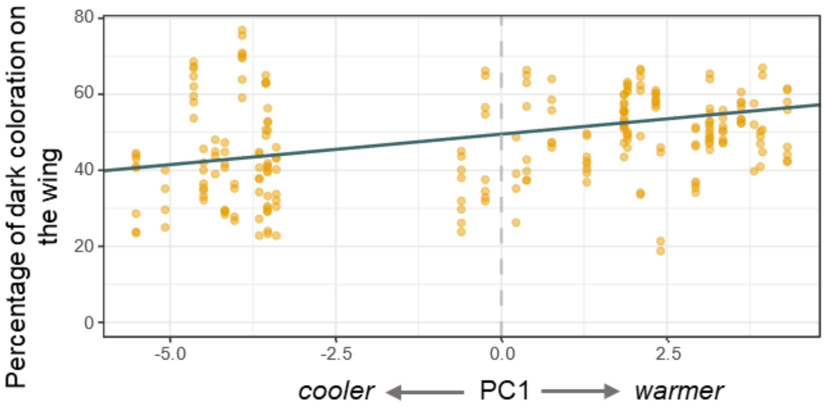


**Figure S7**. Proportion of dark coloration on the surface of butterfly wings among transparent species only as a function of climatic principal component PC1. The relationship is significant and the solid line is drawn from the prediction of the Bayesian phylogenetic mixed model (see Table S15) (N = 235).

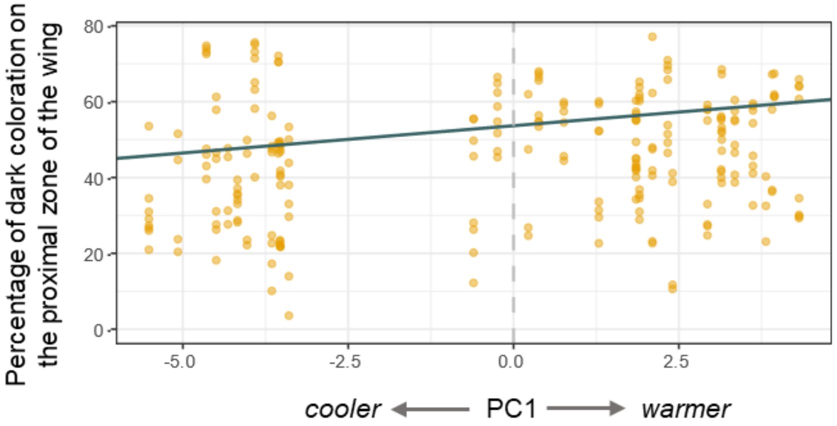


**Figure S8.** Proportion of dark coloration on the surface of proximal zones of butterfly wings among transparent species as a function of climatic principal component PC1. The relationship is significant The solid line is drawn from the prediction of the Bayesian phylogenetic mixed model (see Table S16) (N = 235).

**Table S1.** List of the species and the number of specimens per species studied. All specimens are females. The mean altitude of occurrence of each species is computed based on published data (1), and the mean transmission over the UV-Visible range (300 to 700 nm) of the forewings and hindwings spots for each species are shown. The threshold for the division of the species in two groups - opaque or transparent- was set at 35% of transmission for both wings.

| Genus, species | Number of specimens (females) | Mean altitude of occurrence in meters | Mean transmission % of the Forewing spot in the UV-Visible | Mean transmission % of the Hindwing spot in the UV-Visible | Category of the species |
|---|---|---|---|---|---|
| *Athesis acrisione* | 2 | 752.00 | 63.14 | 57.70 | Transparent |
| *Callithomia lenea* | 4 | 601.37 | 55.34 | 51.31 | Transparent |
| *Ceratinia poecila* | 5 | 768.17 | 31.31 | 27.29 | Opaque |
| *Dircenna dero* | 4 | 515.31 | 67.46 | 58.86 | Transparent |
| *Dircenna loreta* | 3 | 780.39 | 66.04 | 59.21 | Transparent |
| *Dircenna olyras* | 2 | 894.08 | 32.27 | 27.27 | Opaque |
| *Godyris duillia* | 4 | 1404.10 | 73.00 | 71.68 | Transparent |
| *Godyris hewitsoni* | 4 | 2041.00 | 66.75 | 62.41 | Transparent |
| *Godyris panthyale* | 4 | 2023.83 | 80.80 | 71.00 | Transparent |
| *Godyris zavaleta* | 4 | 367.73 | 62.69 | 58.12 | Transparent |
| *Greta morgane* | 2 | 750.07 | 74.59 | 78.45 | Transparent |
| *Heterosais nephele* | 4 | 551.40 | 65.87 | 67.07 | Transparent |
| *Hyalenna paradoxa* | 2 | 2195.87 | 68.64 | 59.61 | Transparent |
| *Hyalyris coeno* | 3 | 831.05 | 26.57 | 1.96 | Opaque |
| *Hyalyris mestra* | 3 | 1447.95 | 61.78 | 66.41 | Transparent |
| *Hyalyris oulita* | 4 | 1208.42 | 29.53 | 13.61 | Opaque |
| *Hypoleria xenophis* | 4 | 710.00 | 60.13 | 61.87 | Transparent |
| *Hypomenitis alphesiboea* | 4 | 1545.38 | 60.24 | 70.71 | Transparent |
| *Hypomenitis ortygia* | 4 | 2030.49 | 71.43 | 59.42 | Transparent |
| *Hypomenitis theudelinda* | 4 | 2166.95 | 63.62 | 66.13 | Transparent |
| *Hypothyris cantobrica* | 4 | 740.47 | 15.10 | 12.86 | Opaque |
| *Ithomia avella* | 4 | 2006.97 | 57.29 | 57.08 | Transparent |
| *Ithomia xenos* | 2 | 1319.83 | 36.96 | 41.94 | Transparent |
| *Mechanitis polymnia* | 5 | 596.00 | 13.35 | 7.99 | Opaque |
| *Melinaea menophilus* | 4 | 846.00 | 4.76 | 4.58 | Opaque |
| *Methona confusa* | 8 | 896.78 | 53.25 | 47.36 | Transparent |
| *Methona curvifascia* | 4 | 416.85 | 52.81 | 48.14 | Transparent |
| *Napeogenes larilla* | 4 | 2146.11 | 62.37 | 63.30 | Transparent |
| *Napeogenes sylphis* | 5 | 411.50 | 43.99 | 73.55 | Transparent |
| *Oleria athalina* | 4 | 2201.62 | 37.95 | 69.26 | Transparent |
| *Oleria baizana* | 4 | 2119.44 | 40.46 | 57.80 | Transparent |
| *Ollantaya olerioides* | 3 | 2029.27 | 57.74 | 65.48 | Transparent |
| *Pagyris cymothoe* | 4 | 1191.62 | 72.16 | 72.91 | Transparent |

| | | | | | |
|---|---|---|---|---|---|
| *Patricia dercyllidas* | 2 | 2058.44 | 47.89 | 56.98 | Transparent |
| *Pteronymia oneida* | 4 | 2179.98 | 52.71 | 61.71 | Transparent |
| *Pteronymia sao* | 2 | 104.86 | 52.91 | 55.21 | Transparent |
| *Pteronymia vestilla* | 4 | 333.96 | 58.99 | 67.56 | Transparent |
| *Scada reckia* | 2 | 743.90 | 51.94 | 43.31 | Transparent |
| *Thyridia psidii* | 4 | 788.93 | 48.52 | 48.36 | Transparent |
| *Tithorea harmonia* | 7 | 543.08 | 3.78 | 2.16 | Opaque |
| *Tithorea tarricina* | 2 | 1285.44 | 2.09 | 6.92 | Opaque |
| *Veladyris pardalis* | 4 | 2036.77 | 59.91 | 38.54 | Transparent |

**Table S2**. Retained Bayesian mixed models correcting for phylogenetic relatedness of the optical relationship between absorption and transmission spectra of the wing spots at different ranges, after backward selection of fixed parameters based on Bayesian P-value criteria. Fixed effect estimates with 95% confidence intervals excluding zero are indicated in bold, associated with a Bayesian P-value (*P < 0.05; **P < 0.01; ***P < 0.001). The factors for which the parameters were not significant are listed below the model.

| **Dependent variable** | **Parameter** | **Posterior mean** | **Lower 95 % Confidence Interval** | **Upper 95 % Confidence Interval** | **Effective size** | **Bayesian P-value** |
|---|---|---|---|---|---|---|
| **Mean absorption of the wing spot over 300 to 2500 nm** | (Intercept) | 13.992 | 11.873 | 16.081 | 156309.039 | < 0.001 *** |
| | **Mean transmission over 300 to 700 nm** | **-8.450** | **-9.124** | **-7.790** | **102746.658** | **< 0.001 *** ** |
| | phylogenetic variance | 5.953 | 0.000 | 10.327 | 5927.130 | |
| | Species identity | 0.385 | 0.000 | 2.097 | 1548.627 | |
| | residual variance | 7.621 | 6.380 | 8.975 | 183185.405 | |
| **Mean absorption of the wing spot over 700 to 2500 nm** | (Intercept) | 6.964 | 4.433 | 9.423 | 361960.677 | < 0.001 *** |
| | **Mean transmission over 300 to 700 nm** | **-5.326** | **-6.091** | **-4.540** | **108436.277** | **< 0.001*** ** |
| | phylogenetic variance | 8.348 | 0.000 | 14.321 | 7731.567 | |
| | Species identity | 0.490 | 0.000 | 2.689 | 1436.113 | |
| | residual variance | 10.727 | 8.941 | 12.581 | 353377.654 | |
| **Mean absorption of the wing spot over 700 to 2500 nm** | (Intercept) | 6.038 | 2.134 | 10.106 | 32631.740 | < 0.01** |
| | **Mean absorption over 300 to 700 nm** | **5.146** | **3.146** | **7.061** | **31417.877** | **< 0.001*** ** |
| | phylogenetic variance | 8.328 | 3.367 | 14.214 | 4269.801 | |
| | Species identity | 0.239 | 0.000 | 1.188 | 3244.589 | |
| | residual variance | 10.096 | 8.402 | 11.851 | 32559.210 | |
| | Other fixed parameters not retained : Type of wing (opaque > transparent); Mean absorption over 300 to 700 nm * Type of wing | | | | | |

**Table S3.** Bayesian mixed models correcting for phylogenetic relatedness of the relationship between the thermal properties of the wing spots and their optical properties at different ranges. Fixed effect estimates with 95% confidence intervals excluding zero are indicated in bold, associated with a Bayesian P-value (*P < 0.05; **P < 0.01; ***P < 0.001).

| Dependent variable | Parameter | Posterior mean | Lower 95 % Confidence Interval | Upper 95 % Confidence Interval | Effective size | Bayesian P-value |
|---|---|---|---|---|---|---|
| Excess temperature ΔT of the wing spot | (Intercept) | 11.932 | 10.033 | 13.761 | 19500.000 | < 0.001 *** |
| | **Mean transmission over 300 to 700 nm** | **-1.546** | **-1.919** | **-1.167** | **13067.170** | **< 0.001 *** ** |
| | phylogenetic variance | 5.232 | 2.449 | 8.410 | 3153.368 | |
| | Species identity | 0.128 | 0.000 | 0.597 | 1979.854 | |
| | residual variance | 1.586 | 1.331 | 1.877 | 18984.158 | |
| | (Intercept) | 12.364 | 10.100 | 14.534 | 18828.512 | < 0.001 *** |
| | **Mean absorption over 300 to 2500 nm** | **0.983** | **0.671** | **1.280** | **14300.000** | **< 0.001 *** ** |
| | phylogenetic variance | 7.256 | 3.170 | 11.700 | 2900.000 | |
| | Species identity | 0.267 | 0.000 | 1.190 | 1860.000 | |
| | residual variance | 1.612 | 1.350 | 1.901 | 19060.100 | |
| | (Intercept) | 12.548 | 10.000 | 14.954 | 19500.000 | < 0.001 *** |
| | **Mean absorption over 700 to 2500 nm** | **0.596** | **0.351** | **0.864** | **14643.765** | **< 0.001 *** ** |
| | phylogenetic variance | 8.961 | 3.520 | 14.847 | 1935.934 | |
| | Species identity | 0.537 | 0.000 | 1.996 | 1608.264 | |
| | residual variance | 1.658 | 1.390 | 1.958 | 17949.578 | |

**Table S4.** Bayesian mixed model correcting for phylogenetic relatedness of the relationship between the mean excess temperature $\Delta T$ of the Dark patches DP of butterfly and the Lighter LP and Transparent patches TP, and the zone of the wing, proximal P or distal D. Fixed effect estimates with 95% confidence intervals excluding zero are indicated in bold, associated with a Bayesian P-value (*P < 0.05; **P < 0.01; ***P < 0.001).

| **Dependent variable** | **Parameter** | **Posterior mean** | **Lower 95 % Confidence Interval** | **Upper 95 % Confidence Interval** | **Effective size** | **Bayesian P-value** |
|---|---|---|---|---|---|---|
| Excess temperature $\Delta T$ | (Intercept) | 20.212 | 18.400 | 22.076 | 591933.290 | < 0.001 *** |
| | **Patch color DP>(LP, TP))** | **11.775** | **12.000** | **11.507** | **592594.820** | **< 0.001 *** ** |
| | **Zone (P>D)** | **-1.253** | **-1.520** | **-0.990** | **596000.000** | **< 0.001 *** ** |
| | **Patch color DP>(LP, TP) x Zone (P>D)** | **-1.290** | **-0.915** | **-1.667** | **592931.730** | **< 0.001 *** ** |
| | phylogenetic variance | 5.170 | 2.750 | 7.975 | 62364.570 | |
| | Species identity | 0.109 | 0.000 | 0.485 | 35697.100 | |
| | residual variance | 8.381 | 8.000 | 8.769 | 593640.220 | |

**Table S5.** Bayesian mixed model correcting for phylogenetic relatedness of the relationship between the mean excess temperature ΔT of patches and patch color (Light patches LP and Transparent patches TP), and wing zone (proximal P or distal D). Fixed effect estimates with 95% confidence intervals excluding zero are indicated in bold, associated with a Bayesian P-value (*P < 0.05; **P < 0.01; ***P < 0.001).

| Dependent variable | Parameter | Posterior mean | Lower 95 % Confidence Interval | Upper 95 % Confidence Interval | Effective size | Bayesian P-value |
|---|---|---|---|---|---|---|
| Excess temperature ΔT | (Intercept) | 10.067 | 8.630 | 11.477 | 45300.000 | < 0.001 *** |
| | **Patch color (LP>TP)** | **2.813** | **1.609** | **4.080** | **9450.000** | **< 0.001 *** ** |
| | **Zone (P>D)** | **1.683** | **2.020** | **1.340** | **49500.000** | **< 0.001 *** ** |
| | **Patch color (LP>TP) x Zone (P>D)** | **2.055** | **2.433** | **1.670** | **49500.000** | **< 0.001 *** ** |
| | phylogenetic variance | 1.979 | 0.368 | 4.029 | 2602.930 | |
| | Species identity | 0.817 | 0.000 | 1.624 | 4488.541 | |
| | residual variance | 2.925 | 2.733 | 3.118 | 49500.000 | |

**Table S6**. Bayesian mixed model correcting for phylogenetic relatedness of the relationship between the mean temperature excess ΔT and Dark patches DP vs Dark Veins DV, and wing zone (proximal P or distal D). Fixed effect estimates with 95% confidence intervals excluding zero are indicated in bold, associated with a Bayesian P-value (*P < 0.05; **P < 0.01; ***P < 0.001).

| **Dependent variable** | **Parameter** | **Posterior mean** | **Lower 95 % Confidence Interval** | **Upper 95 % Confidence Interval** | **Effective size** | **Bayesian P-value** |
|---|---|---|---|---|---|---|
| Excess temperature ΔT | (Intercept) | 19.570 | 17.441 | 21.620 | 48100.00 | < 0.001 *** |
| | **Patch color (DP>DV)** | **5.517** | **5.870** | **5.163** | **49500.00** | **< 0.001 *** ** |
| | **Zone (P>D)** | **-1.255** | **-1.610** | **-0.918** | **49500.00** | **< 0.001 *** ** |
| | **Patch color (DV>DP) x Zone (P>D)** | **-3.491** | **-3.010** | **-3.958** | **49500.00** | **< 0.001 *** ** |
| | phylogenetic variance | 5.896 | 1.310 | 11.043 | 3380.00 | |
| | Species identity | 1.567 | 0.000 | 3.357 | 5550.00 | |
| | residual variance | 14.068 | 13.500 | 14.706 | 49500.00 | |

**Table S7**. Retained Bayesian mixed model correcting for phylogenetic relatedness of the relationship between the mean temperature excess ΔT and light patches LP vs the Light Veins LV, and wing zone (proximal P or distal D), after backward selection of fixed parameters based on Bayesian P-value criteria. Fixed effect estimates with 95% confidence intervals excluding zero are indicated in bold, associated with a Bayesian P-value (*P < 0.05; **P < 0.01; ***P < 0.001). The factors for which the parameters were not significant are listed below the model.

| Dependent variable | Parameter | Posterior mean | Lower 95 % Confidence Interval | Upper 95 % Confidence Interval | Effective size | Bayesian P-value |
|---|---|---|---|---|---|---|
| Excess temperature ΔT | (Intercept) | 8.514984 | 6.030 | 11.142413 | 91944.511 | < 0.001 *** |
| | **Patch color (LV> LP)** | **3.513077** | **3.050** | **3.980312** | **464070.148** | **< 0.001 *** ** |
| | **Zone (P>D)** | **1.33338** | **0.947** | **1.710718** | **596000** | **< 0.001 *** ** |
| | phylogenetic variance | 7.346833 | 0.000 | 15.386159 | 4531.189 | |
| | Species identity | 1.560804 | 0.000 | 7.593318 | 1492.881 | |
| | residual variance | 5.784633 | 5.190 | 6.395479 | 596000 | |
| | Other fixed parameters not retained in the best model : Patch color (LV> LP) x Zone (P>D) | | | | | |

**Table S8.** Coordinates of WorldClim variables on the climatic principal component PC1.

| WorldClim Variables | Coordinates on PC1 |
|---|---|
| Min Temperature of Coldest Month | 0.937 |
| Mean Temperature of Driest Quarter | 0.926 |
| Mean Temperature of Warmest Quarter | 0.915 |
| Annual Mean Temperature | 0.908 |
| Mean Temperature of Wettest Quarter | 0.901 |
| Mean Temperature of Coldest Quarter | 0.895 |
| Max Temperature of Warmest Month | 0.885 |
| Precipitation of Warmest Quarter | 0.858 |
| Precipitation of Wettest Quarter | 0.776 |
| Annual Precipitation | 0.757 |
| Precipitation of Wettest Month | 0.759 |
| Isothermality (Mean Diurnal Range/Temperature Annual Range) (×100) | -0.549 |
| Precipitation of Driest Quarter | 0.457 |
| Temperature Seasonality (standard deviation ×100) | 0.432 |
| Precipitation of Driest Month | 0.443 |
| Mean Diurnal Range (Mean of monthly (max temp - min temp)) | -0.417 |
| Precipitation of Coldest Quarter | 0.337 |
| Precipitation Seasonality (Coefficient of Variation) | 0.107 |
| Temperature Annual Range (Max Temperature of Warmest Month - Min Temperature of Coldest Month) | 0.011 |

**Table S9**. Coordinates of WorldClim variables on the climatic principal component PC2.

| WorldClim Variables | Coordinates on PC2 |
|---|---|
| Precipitation of Coldest Quarter | 0.907 |
| Temperature Annual Range (Max Temperature of Warmest Month - Min Temperature of Coldest Month) | -0.833 |
| Precipitation of Driest Quarter | 0.812 |
| Precipitation of Driest Month | 0.802 |
| Annual Precipitation | 0.608 |
| Precipitation Seasonality (Coefficient of Variation) | -0.607 |
| Mean Diurnal Range (Mean of monthly (max temp - min temp)) | -0.537 |
| Isothermality (Mean Diurnal Range/Temperature Annual Range) (×100) | 0.507 |
| Temperature Seasonality (standard deviation ×100) | -0.452 |
| Precipitation of Wettest Quarter | 0.373 |
| Max Temperature of Warmest Month | -0.398 |
| Mean Temperature of Wettest Quarter | -0.367 |
| Precipitation of Wettest Month | 0.322 |
| Mean Temperature of Warmest Quarter | -0.344 |
| Annual Mean Temperature | -0.322 |
| Mean Temperature of Coldest Quarter | -0.291 |
| Precipitation of Warmest Quarter | 0.259 |
| Mean Temperature of Driest Quarter | -0.237 |
| Min Temperature of Coldest Month | -0.165 |

**Table S10.** Retained Bayesian mixed model correcting for phylogenetic relatedness of the relationship between the mean transmission of the wing spots over the range 300 to 700 nm and climatic variables, after backward selection of fixed parameters based on Bayesian P-value criteria. Fixed effect estimates with 95% confidence intervals excluding zero are indicated in bold, associated with a Bayesian P-value (*P < 0.05; **P < 0.01; ***P < 0.001). The factors for which the parameters were not significant are listed below the model.

| Dependent variable | Parameter | Posterior mean | Lower 95 % Confidence Interval | Upper 95 % Confidence Interval | Effective size | Bayesian P-value |
|---|---|---|---|---|---|---|
| Mean transmission of the wing spot over 300 to 700 nm | (Intercept) | 38.144 | 22.690 | 53.214 | 9323.783 | < 0.001*** |
| | **PC2** | **4.698** | **0.202** | **9.504** | **19500.000** | **0.048 *** |
| | phylogenetic variance | 291.857 | 63.653 | 570.576 | 1433.784 | |
| | Species identity | 144.982 | 35.479 | 275.447 | 1282.366 | |
| | residual variance | 60.095 | 49.943 | 70.440 | 19500.000 | |
| | Other fixed parameters not retained: PC1; PC1 x PC2 | | | | | |

**Table S11**. Retained Bayesian mixed model correcting for phylogenetic relatedness of the relationship between the mean absorption of the wing spots over the range 300 to 2500 m and the climatic variables, after backward selection of fixed parameters based on Bayesian P-value criteria. Fixed effect estimates with 95% confidence intervals excluding zero are indicated in bold, associated with a Bayesian P-value (*P < 0.05; **P < 0.01; ***P < 0.001). The factors for which the parameters were not significant are listed below the model.

| **Dependent variable** | **Parameter** | **Posterior mean** | **Lower 95 % Confidence Interval** | **Upper 95 % Confidence Interval** | **Effective size** | **Bayesian P-value** |
|---|---|---|---|---|---|---|
| Mean absorption of the wing spot over 300 to 2500 nm | (Intercept) | 18.187 | 12.226 | 24.068 | 14519.316 | < 0.001*** |
| | phylogenetic variance | 45.574 | 12.876 | 87.789 | 1597.004 | |
| | Species identity | 14.485 | 0.001 | 30.014 | 2264.057 | |
| | residual variance | 18.862 | 15.813 | 22.190 | 19500.000 | |
| | Fixed parameters not retained: PC1; PC2; PC1 x PC2 | | | | | |

**Table S12**. Retained Bayesian mixed model correcting for phylogenetic relatedness of the relationship between the mean absorption of the wing spots over the range 700 to 2500 m and the climatic variables, after backward selection of fixed parameters based on Bayesian P-value criteria. Fixed effect estimates with 95% confidence intervals excluding zero are indicated in bold, associated with a Bayesian P-value (*P < 0.05; **P < 0.01; ***P < 0.001). The factors for which the parameters were not significant are listed below the model.

| **Dependent variable** | **Parameter** | **Posterior mean** | **Lower 95 % Confidence Interval** | **Upper 95 % Confidence Interval** | **Effective size** | **Bayesian P-value** |
|---|---|---|---|---|---|---|
| Mean absorption of the wing spot over 700 to 2500 nm | (Intercept) | 9.835 | 5.645 | 14.080 | 18746.162 | < 0.001*** |
| | phylogenetic variance | 25.140 | 10.090 | 43.846 | 1973.388 | |
| | Species identity | 2.197 | 0.000 | 7.761 | 1023.411 | |
| | residual variance | 16.082 | 13.416 | 18.877 | 15939.104 | |
| | Fixed parameters not retained: PC1; PC2; PC1 x PC2 | | | | | |

**Table S13.** Retained Bayesian mixed model correcting for phylogenetic relatedness of the relationship between the mean temperature excess ΔT of the full wing and the climatic variables, after backward selection of fixed parameters based on Bayesian P-value criteria. Fixed effect estimates with 95% confidence intervals excluding zero are indicated in bold, associated with a Bayesian P-value (*P < 0.05; **P < 0.01; ***P < 0.001). The factors for which the parameters were not significant are listed below the model.

| **Dependent variable** | **Parameter** | **Posterior mean** | **Lower 95 % Confidence Interval** | **Upper 95 % Confidence Interval** | **Effective size** | **Bayesian P-value** |
|---|---|---|---|---|---|---|
| Mean excess temperature ΔT of the full wing | (Intercept) | 13.317 | 11.100 | 15.647 | 18800.000 | < 0.001*** |
| | **PC1** | **0.810** | **0.198** | **1.421** | **17912.730** | **< 0.05*** |
| | Phylogenetic variance | 7.464 | 2.770 | 12.917 | 1750.074 | |
| | Species identity | 0.828 | 0.000 | 2.324 | 1816.949 | |
| | Residual variance | 2.256 | 1.890 | 2.658 | 18079.378 | |
| | Other fixed parameters not retained: PC2; PC1 x PC2; category of the species (transparent or opaque); area of the wing; identity of the wing (forewing or hindwing) | | | | | |

**Table S14.** Retained Bayesian mixed model correcting for phylogenetic relatedness of the relationship between the proportion of dark patches on the surface of each wing and climatic variables, after backward selection of fixed parameters based on Bayesian P-value criteria. Fixed effect estimates with 95% confidence intervals excluding zero are indicated in bold, associated with a Bayesian P-value (*P < 0.05; **P < 0.01; ***P < 0.001). The factors for which the parameters were not significant are listed below the model.

| **Dependent variable** | **Parameter** | **Posterior mean** | **Lower 95 % Confidence Interval** | **Upper 95 % Confidence Interval** | **Effective size** | **Bayesian P-value** |
|---|---|---|---|---|---|---|
| Proportion of dark patches on the wings | (Intercept) | 50.492 | 40.770 | 59.900 | 48300 | < 0.001 *** |
| | **PC1** | **4.558** | **1.599** | **7.530** | **11300** | **< 0.01 **** |
| | **Wing identity Forewing>Hindwing** | **-3.886** | **-6.079** | **-1.731** | **49900** | **< 0.001 ***** |
| | phylogenetic variance | 123.971 | 27.181 | 242.779 | 2070 | |
| | Species identity | 22.540 | 0.000 | 68.837 | 1270 | |
| | Residual variance | 92.840 | 77.400 | 109.543 | 29163.515 | |
| | Other fixed parameters not retained: PC2; PC1 x PC2; category of the species (transparent or opaque); area of the wing | | | | | |

**Table S15.** Retained Bayesian mixed model correcting for phylogenetic relatedness of the relationship between the proportion of dark patches on the surface of each wing among transparent species as a function of climatic variables, after backward selection of fixed parameters based on Bayesian P-value criteria. Fixed effect estimates with 95% confidence intervals excluding zero are indicated in bold, associated with a Bayesian P-value (*P < 0.05; **P < 0.01; ***P < 0.001). The factors for which the parameters were not significant are listed below the model.

| **Dependent variable** | **Parameter** | **Posterior mean** | **Lower 95 % Confidence Interval** | **Upper 95 % Confidence Interval** | **Effective size** | **Bayesian P-value** |
|---|---|---|---|---|---|---|
| Proportion of dark patches on transparent wings | (Intercept) | 49.073 | 39.100 | 58.651 | 49000.000 | < 0.001 *** |
| | **PC1** | **5.219** | **2.780** | **7.659** | **38812.461** | **< 0.001 *** ** |
| | Phylogenetic variance | 104.112 | 44.500 | 180.245 | 5872.326 | |
| | Species identity | 2.086 | 0.000 | 11.100 | 3568.089 | |
| | Residual variance | 65.149 | 53.200 | 78.500 | 45847.698 | |
| | Other fixed parameters not retained: PC2; PC1 x PC2; area of the wing; identity of the wing (forewing or hindwing) | | | | | |

**Table S16**. Retained Bayesian mixed model correcting for phylogenetic relatedness of the relationship between the proportion of colored patches in the proximal zone of each wing among transparent species as a function of climatic variables, after backward selection of fixed parameters based on Bayesian P-value criteria. Fixed effect estimates with 95% confidence intervals excluding zero are indicated in bold, associated with a Bayesian P-value (*P < 0.05; **P < 0.01; ***P < 0.001). The factor for which the parameters were not significant is listed below the model.

| Dependent variable | Parameter | Posterior mean | Lower 95 % Confidence Interval | Upper 95 % Confidence Interval | Effective size | Bayesian P-value |
|---|---|---|---|---|---|---|
| Proportion of dark patches in the proximal zone of transparent wings | (Intercept) | 53.271 | 42.800 | 63.465 | 1280000 | < 0.001 *** |
| | **PC1** | **4.676** | **1.490** | **7.851** | **30600** | **< 0.001 *** |
| | **PC2** | **3.292** | **0.641** | **5.967** | **984000** | **< 0.05*** |
| | **Wing identity forewing > hindwing** | **-16.474** | **-18.900** | **-14.054** | **1380000** | **< 0.001 *** |
| | Phylogenetic variance | 104.876 | 0.000 | 206.986 | 10000 | |
| | Species identity | 17.428 | 0.000 | 81.513 | 3413.408 | |
| | Residual variance | 89.498 | 72.400 | 107.592 | 1092255.180 | |
| | Other fixed parameters not retained: PC1 x PC2 | | | | | |

**Table S17**. Retained Bayesian mixed model correcting for phylogenetic relatedness of the relationship between the total area of the hindwing plus forewing of each butterfly and the climatic variables. Retained Bayesian mixed model corrected for phylogenetic relatedness, after backward selection of fixed parameters based on Bayesian P-value criteria. Fixed effect estimates with 95% confidence intervals excluding zero are indicated in bold, associated with a Bayesian P-value (*P < 0.05; **P < 0.01; ***P < 0.001). The factor for which the parameters were not significant is listed below the model.

| Dependent variable | Parameter | Posterior mean | Lower 95 % Confidence Interval | Upper 95 % Confidence Interval | Effective size | Bayesian P-value |
|---|---|---|---|---|---|---|
| Total surface of the hindwing plus forewing of each butterfly | (Intercept) | 759.086 | 519.548 | 997.923 | 59383.942 | < 0.001*** |
| | phylogenetic variance | 86397.084 | 44978.920 | 135603.575 | 4361.406 | |
| | Species identity | 1206.417 | 0.000 | 7524.155 | 1223.185 | |
| | residual variance | 5080.964 | 3795.435 | 6458.049 | 59000.000 | |
| | Fixed parameters not retained: PC1; PC2; PC1 x PC2 | | | | | |
